# Mitochondrial fitness influences neuronal excitability of dopaminergic neurons from patients with idiopathic form of Parkinson’s disease

**DOI:** 10.1101/2023.04.28.538698

**Authors:** Paula Chlebanowska, Agata Szlaga, Anna Tejchman-Skrzyszewska, Marta Kot, Pawel Konieczny, Klaudia Skrzypek, Agata Muszynska, Malgorzata Sobocinska, Krystyna Golebiowska, Pawel Labaj, Anna Blasiak, Marcin Majka

**Affiliations:** Jagiellonian University Medical College, Faculty of Medicine, Institute of Pediatrics, Department of Transplantation, Krakow, Poland; Department of Neurophysiology and Chronobiology, Institute of Zoology and Biomedical Research, Faculty of Biology, Jagiellonian University, Krakow, 30-387, Poland; Bioinformatics Research Group, Malopolska Centre of Biotechnology, Jagiellonian University, Gronostajowa 7A, 30-387 Kraków, Poland; Department of Pharmacology, Institute of Pharmacology, Polish Academy of Sciences, 12 Smetna, 31-343, Kraków, Poland.

**Author notes:** Corresponding authors: Prof. Marcin Majka, PhD, DSc, Jagiellonian University Medical College, Faculty of Medicine, Institute of Pediatrics, Department of Transplantation, Wielicka 265, 30-663 Krakow, Poland, Prof. Anna Blasiak, PhD, DSc, Department of Neurophysiology and Chronobiology, Institute of Zoology and Biomedical Research, Faculty of Biology, Jagiellonian University, Krakow, 30-387, Poland. These authors contributed equally to this work (Paula Chlebanowska, Agata Szlaga). Co-seniors authors (Anna Blasiak, Marcin Majka).

**Keywords:** Parkinson’s disease, dopaminergic neurons, mitochondria, iPS

## Abstract

Parkinson disease is the second most common neurodegenerative disease defined by presence of Lewy bodies and the loss of dopaminergic neurons in the substantia nigra pars compacta (SNc). There are three types of PD - familial, early-onset and idiopathic. Idiopathic PD (IPD) accounts for approximately 90% of all PD cases. Mitochondrial dysfunction accompanies the pathogenesis of Parkinson’s disease. Loss of mitochondrial function increases oxidative stress and calcium buffering, which in turn hinders the production of ATP and disrupts the functioning of dopaminergic neurons. The main barrier in PD research was the lack of proper human models to study the mechanisms of PD development and progression. Using induced pluripotent stem (iPS) cells we generated patient-specific dopaminergic neurons.

We observed differences in the mitochondria fitness but not differences in mitochondria mass, morphology or membrane potential. Expression of OXPHOS mitochondrial complexes were lower in PD patients in comparison to control group what resulted in changes in mitochondria respiratory status. We observed also lower expression levels of Na^+^/K^+^-ATPase subunits and ATP-sensitive K^+^ (K-ATP) channel subunits. The lower oxygen consumption rate and extracellular acidification rate values were observed in dopaminergic progenitors and iPSC from PD patients compared to the control group. Importantly, observed decrease in the availability of ATP and in the energy consumption, as well as changes in acidification, may constitute contributing factors to the observed reduced neuronal excitability of PD patients dopaminergic neurons.

## Introduction

Neurodegenerative diseases (e.g., Alzheimer’s Disease, Parkinson’s Disease, a Amyotrophic Lateral Sclerosis) are becoming the growing medical problems for conventional medicine. The frequency of neurodegenerative disease prevalence rises due to aging society. The increasing number of people suffering from these disorders causes an economic challenge for our society ^1, 2^. Those patients are very demanding as far as the proper care is concerned. They need constant attention and the presence of family members or qualified medical help. This creates an increased burden on the health care systems around the world and calls for new treatments to be found.

Parkinson’s disease (PD) is the second most common neurodegenerative disease defined by presence of Lewy bodies and the loss of dopaminergic neurons in the substantia nigra pars compacta (SNc) ^3^. PD manifests itself by many clinical symptoms such as e.g. motor bradykinesia, rigidity, tremor and non-motor apathy, memory complaints, sleep problems, mood disturbances, fatigue ^4^. There are three types of PD - familial, early-onset and idiopathic. The familial form of PD is identified in 5 to 10% of the cases. The early-onset PD or young – onset PD (YOPD) is a rare type of PD affecting people between the age of 20 to 50 years and is observed in around 7.5% of PD patients ^5^. Idiopathic PD (IPD) is the most common type of disease ^6^ and accounts for approximately 90% of all PD cases ^6, 7^. In the case of YOPD and IPD the exact etiology and pathomechanisms responsible for PD are not known, however, the genetic components, neuroinflammation, environmental influence or lysosomal defect are suggested as the major causes of PD ^8–10^.

Mitochondrial dysfunction accompanies the pathogenesis of many neurodegenerative diseases, including Parkinson’s disease, in which dopaminergic neurons gradually degenerate ^10, 11^. Loss of mitochondrial function increases oxidative stress and calcium buffering, which in turn hinders the production of ATP and disrupts the functioning of dopaminergic neurons^10^.

Until recently, the main barrier in PD research was the lack of proper human models to study the mechanisms of PD development and progression. A significant new tool to study mechanisms and causes of PD are induced pluripotent stem (iPS) cells ^12^. “Patient-specific” cell-based models can reproduce molecular background of particular case, without ethical concerns ^13, 14^. Moreover, the iPS cells technology gives us the opportunity to generate dopaminergic neurons and to study the etiology of PD ^15^.

Here, we created and characterized iPS cell lines from healthy volunteers and patients with IPD and used this model to identify the drivers responsible for IPD.

## Materials and methods

### Cell Culture

Peripheral Blood Mononuclear Cells (PBMCs) were cultured in medium QBSF-60 (VWR, Radnor, PA, USA) contains 10 µL/mL ascorbic acid, 100 U/ml Penicillin/Streptomycin and growth factors: 1,5 µM dexamethasone, 2 U/ml EPO, 10 ng/ml IL-3, 50 ng/ml SCF, 40 ng/ml IGF-1 (all supplements from Peprotech, London, United Kingdom). Medium with all supplements is called Expansion Medium.

Mouse embryonic fibroblasts (MEFs) were cultured in Dulbecco’s Modified Eagle’s Medium (DMEM) with 4,5 g/L glucose (PAN Biotech, Aidenbach, Germany) and with supplement 2 mM L-glutamine, 100 U/ml Penicillin/Streptomycin (both from Thermo Fisher Scientific, Waltham, MA, USA) and 10% v/v FBS (Eurx, Gdansk, Poland). MEFs were inactivated by 3h incubation cells with mitomycin C (Sigma–Aldrich, Saint Louis, MO, USA) diluted in medium.

Induced Pluripotent Stem (iPS) cells were generated from PBMC. The cells were cultured on Matrigel (feeder free) (Corning, New York, NY, USA) and feeder layer of iMEF. The iPS cells were cultured in serum free iPS medium, based on DMEM/F12 with 2mM Glutamax, supplemented with 20% KSR, 100 µM Non-Essential Amino Acids, 100 U/ml Penicillin/Streptomycin, 10 ng/ml bFGF (all from Thermo Fisher Scientific) and 100 µM β-mercaptoethanol (Sigma–Aldrich). The iPS medium for feeder free culture was conditioned by 24h incubation on iMEF. For both culture conditions of iPS cells the medium was changed every day. iPS cells were passaged every 3-4 days with Accutase (Lonza, Basel, Switzerland) and seeded on new plates with density 1:5 – 1:10 in iPS medium with ROCK inhibitor Y-27632 (Sigma-Aldrich). Before freezing the cells were incubated with ROCK inhibitor Y-27632 (Sigma-Aldrich) by 1h. The iPS freezing medium was consisted with 90% FBS (Eurx), 10% DMSO (Sigma-Aldrich), 10 µM Y-27632 (Sigma–Aldrich). iPS cells were cryopreserved in liquid nitrogen in density of 1 × 10^6^ cells/ml. The cells were cultured at 37 °C in a humidified atmosphere of 5% CO_2_.

### Isolation of PBMC from full blood

The study was approved by the Jagiellonian University Bioethical Committee in Krakow (decision number KBET/173/B/2012). The full blood was drawn from five healthily volunteers and seven PD patients. Before the volunteers were enrolled to the study, each person gave informed consent to participate in the study. PBMCs were isolated by centrifugation on Pancoll (PAN Biotech, Aidenbach, Germany) gradient. 10 ml of full blood was mixed with the same volume of cold PBS (Eurx). This mixture was gently overlaid on 10 ml of cold Pancoll. The tube was centrifuged at room temperature at 1900 rpm for 30 min with the brakes off. The mononuclear cells (buffy coat) from the interface were collected and transferred to the new tube with PBS. The cells were centrifuged at 1300 rpm for 10 min. The supernatant was removed and PBS was added. The cells were washed in the same manner 3 times. Isolated PBMCs were cultured in Expansion Medium for 7 days. Every second day the medium was changed.

### Reprograming of PBMC

Expanded PBMCs were transduced Sendai virus reprogramming vector – according to CytoTune® 2.0 Sendai (Thermo Fisher Scientific). Before cells infection, PBMCs were counted and plated on well of 96 well plate in density 2.5 × 10^⁴^ cells/well. Mixture of three different vectors were added to the well: KOS MOI=5, hc-Myc MOI=5, hKlf4 MOI=3. Cells were incubated overnight. Next day, the virus was washed out by centrifugation. PBMCs were plated on 100 mm plate with iMEF. The medium was changed at day 5 and 7 in proportion (1:1) iPS medium and MEF medium. From day 9, only iPS medium was used. The iPS medium was changed every 2 – 3 days. Between days 21 and 28, the colonies were ready to be picked. About 24 colonies were transferred to the prepared 12 well plates with iMEF (1 colony / 1 well). The iPS colonies were scratched and passaged. The iPS medium was changed every day. After 10 passages the iPS clones with proper morphology were tested and cryopreserved. Five representative clones from healthy volunteers and seven from PD patients were selected to further experiments. Differences between clones from each volunteer were not detected, so representative clones were chosen.

### RNA isolation and reverse transcription

The analysis of gene expression was made by RT-PCR at mRNA level. For total isolation RNA the Universal RNA/miRNA Purification Kit (Eurx) was used. The isolation was made according to the manufacturer’s instruction. After isolation the RNA concentration was measured. The revers transcription (RT) reaction was performed by M-MLV reverse transcription kit (Promega, Madison, WI, USA).

### RT-PCR

To the RT-PCR reactions was used Taq PCR Master Mix (Eurx) and primers with optimal annealing temperature of 55 °C. The presence of Sendai virus genome was checked by primers from Sendai Virus Kit. The primer’s sequences (5’ to 3’): NANOG for: TGAACCTCAGCTACAAACAG, NANOG rev: TGGTGGTAGGAAGAGTAAAG, OCT4 for: ATGGCGGGACACCTGGCTT, OCT4 rev: GGGAGAGCCCAGAGTGGTGACG, TERT for: TGTGCACCAACATCTACAAG, TERT rev: GCGTTCTTGGCTTTCAGGAT, GAPDH for: CAAAGTTGTCATGGATGACC, GAPDH rev: CCATGGAGAAGGCTGGGG.

### RT-qPCR

The method used to analyze gene expression at mRNA level was RT-qPCR. Reactions were performed with Blank qPCR Master Mix 2× (Eurx) and the TaqMan Expression Assays (LMX1A Hs00898455_m1, FOXA2 Hs00232764_m1, NURR1 Hs01117527_g1, TH Hs00165941_m1, TUBB Hs00801390_s1, ABCC8 (Hs01093752_m1); ABCC9 (Hs00245832_m1); KCNJ8 (Hs00958961_m1); KCNJ11 (00265026_s1); ATP1A1 (Hs 00933601_m1); ATP1A2 (Hs00265131_m1); ATP1A3 (Hs 00958036_m1); ATP1A4 (Hs00380134_m1); ATP1B1 (Hs00426868_g1); ATP1B2 (Hs01020302_g1); ATP1B3 (Hs00740857_mH); ATP1B4 (Hs00201320_m1). GAPDH Hs02758991_g1) (Thermo Fisher Scientific) by Quant Studio 7 Flex System (Applied Biosystems, Foster City, CA, USA). The level of analyzed gene expression was normalized to the level of TUBB gene. The method used to analyze was the 2^−ΔΔCt^.

### Immunofluorescent Staining

Cells were washed with PBS and fixed in 4% paraformaldehyde for 20 minutes at room temperature. Then, cells were washed three times and permeabilized in 0,01% Triton X-100 for 5 minutes at room temperature. After incubation, cells were washed by PBS and blocked in 3% bovine serum albumin (BSA, Sigma–Aldrich) for 30 min at room temperature. Subsequently, the primary antibodies diluted in 3% BSA were added to cells. Antibodies such as: mouse anti-tubulin antibody, beta III isoform (Tuj1 MAB1637, Sigma–Aldrich), rabbit anti-tyrosine hydroxylase antibody (TH AB152, Merck-Millipore, CA, USA). After 24 h incubation in 4 °C, cells were washed three times with PBS and incubated with Hoechst (Sigma–Aldrich) and secondary goat anti-rabbit or anti-mouse antibodies by 1h in the dark at room temperature. Secondary antibodies were conjugated with: Alexa Fluor 555 (Thermo Fisher Scientific) or Alexa Fluor 488 (Thermo Fisher Scientific). The secondary antibodies were diluted in 3% BSA. Stained cell were analyzes with fluorescent microscope (Olympus Corporation, Tokyo, Japan).

### Alkaline Phosphatase (AP) Staining

The medium was removed. Cells were washed by PBS and fixed 10 min in 4% paraformaldehyde. After the incubation, cells were washed three times by PBS. Subsequently, cells were incubated in staining solution (1 M Tris-HCl pH 9.5, 5 M NaCl, 1 M MgCl2, NBT-BCIP, H_2_O) (Roche) in the dark for 10 min at room temperature. Then, the solution was removed and cells were washed with PBS. The staining was pictured by microscope IX70 (Olympus Corporation, Tokyo, Japan).

### Teratoma Formation Assay

Approximately 2 x 10^6^ iPS cells were suspended in a 1:1 ratio the PBS and growth factor reduced Matrigel (Corning). Prepared cells were injected into the left dorsal flank of adult female non-obese diabetic / severe combined immunodeficiency (NOD-SCID) mouse. The health of mice and formation of teratomas were checked every day. About two months after injection the tumors were excised. After tumor removal, hematoxylin-eosin (H/E) staining was made. All the experimental protocols were approved the 2nd Local Institutional Animal Care and Use Committee (IACUC) in Krakow (decision numbers 162/2015 and 11/2018).

### Neural differentiation

iPS cells were plated (3,5-4 x 10^4^ cells/cm^2^) on Matrigel at medium KSR: DMEM high glucose, 15% knockout serum replacement, 2 mM L-glutamine, 10 μM βmercaptoethanol and 100 U/ml Penicillin/Streptomycin (all from Thermo Fisher Scientific) with supplements LDN193189 (100nM, Stemgent, Cambridge, MA), SB431542 (10μM, Sigma-Aldrich). Next two days, to the medium from day 0 supplements were added: SHH C25II (100 ng/ml, Peprotech), Purmorphamine (2 μM, Stemgent) and FGF8 (100 ng/ml, Peprotech). On day 3 and 4, to the medium was added CHIR99021 (CHIR; 3 μM, Stemgent). From day 5 the KSR medium was gradually shifted to N2 medium (KSR:N2): day 5 and 6 - 3:1, day 7 and 8 – 1:1, day 9 and 10 – 1:3. N2 medium consisted of DMEM/F12, N2 supplement and 100 U/ml Penicillin/Streptomycin (all from Thermo Fisher Scientific). From day 5 to 10 the medium was supplemented CHIR99021 (CHIR; 3 μM, Stemgent) and LDN193189 (100 nM, Stemgent). On day 5 and 6 SHH C25II (100 ng/ml, Peprotech), Purmorphamine (2 μM, Stemgent) and FGF8 (100 ng/ml, Peprotech) was added to the medium. From day 11 to 19, the medium was consisted of Neurobasal Plus, B27 Plus supplement, BDNF (brain-derived neurotrophic factor, 20 ng/ml; Peprotech), ascorbic acid (AA; 0.2 mM, Sigma-Aldrich), GDNF (glial cell line-derived neurotrophic factor, 20 ng/ml; Peprotech), TGFβ3 (transforming growth factor type β3, 1 ng/ml; Peprotech), dibutyryl cAMP (0.5 mM; Sigma-Aldrich), 2 mM L-glutamine (Thermo Fisher Scientific) and 100 U/ml Penicillin/Streptomycin (Thermo Fisher Scientific). On day 11 and 12, to the medium was added CHIR99021 (CHIR; 3 μM, Stemgent). On day 20, cells were dissociated by Accutase (Lonza) and seeded in high density (3-4 x 10^5^ cells/cm^2^) on earlier coated plate by Polyornithine (PO; 15 μg/ml)/ Laminin (1 μg/ml)/ Fibronectin (2 μg/ml). From day 20, the medium was consisted of Neurobasal Plus, B27 Plus supplement, BDNF (brain-derived neurotrophic factor, 20 ng/ml; Peprotech), ascorbic acid (AA; 0.2 mM, Sigma-Aldrich), GDNF (glial cell line-derived neurotrophic factor, 20 ng/ml; Peprotech), TGFβ3 (transforming growth factor type β3, 1 ng/ml; Peprotech), dibutyryl cAMP (0.5 mM; Sigma-Aldrich), 100 U/ml Penicillin/Streptomycin (Thermo Fisher Scientific) and DAPT (10 nM; Stemgent). The medium was changed every day, until the desired maturation point.

### Extracellular concentration of DA, 5-HT, and Their Metabolites

Extracellular concentration of DA, 5-HT, DOPAC, HVA, and 5-HIAA were measured using a high-performance liquid chromatography (HPLC) with electrochemical detection. Briefly, the samples of medium from over dopaminergic neurons were centrifuged at 10.000 g for 10 min at 4 °C. The supernatants were filtered over Millipore Ultrafree-MC Centrifugal filters with a 0.22 μm pore size hydrophilic PVDF membrane (Millipore) and analyzed by HPLC system. Chromatography was performed using an Ultimate 3000 System (Dionex, USA), electrochemical detector Coulochem III (model 5300; ESA, USA) with a 5020-guard cell, a 5040 amperometric cell and a Hypersil Gold C18 analytical column (3 μm, 100 × 3 mm; Thermo Fisher Scientific, USA). The mobile phase consisted of 0.1 M KH_2_PO_4_ buffer at pH 3.8, 0.5 mM Na_2_EDTA, 100 mg/L 1-octanesulfonic acid sodium salt and 2% methanol. The flow rate during analysis was set at 0.6 mL/min and the applied potential of a guard cell was 600 mV, while that of an amperometric cell was 300 mV with a sensitivity set at 10 nA/V. The chromatographic data were processed by Chromeleon v.6.80 (Dionex) software package.

### TEM imaging

The iPS cells were fixed in a monolayer with 2.5% solution of glutaraldehyde for 3h at room temperature, then scraped with a cell scraper and rinsed in the buffer with sucrose (5.8 g/100 mL). Next the cell pellets were postfixed in buffered 1% osmium tetroxide for 2 h. In the next steps, material was rinsed in the buffer with sucrose (5.8 g/100 mL) and dehydrated in ethanol series (50%, 70%, 90%, 95%, 100%) and acetone, and embedded in epoxy resin Epon 812 (Serva, Heidelberg, Germany). The ultrathin sections (90 nm thick) were doubly contrasted with 2% uranyl acetate and lead citrate and subsequently examined and photographed under a Jeol JEM 2100 transmission electron microscope at 80 kV.

### MitoTracker

To label mitochondria in iPS, cells in monolayer were incubated with 150 nM of MitoTracker™ Deep Red (Invitrogen, M22426) in standard culture medium for 30 min at 37°C in darkness. Mitotracker dye passively diffuse across the plasma membrane and accumulate in active mitochondria. Next the cells were harvested with Accutase solution (BioLegend, USA), rinsed in PBS and analyzed using Attune NxT Software v2.2 on Attune Nxt Flow cytometer (Thermo Fisher Scientific). Unstained cells were used as negative control to determine the background fluorescence and autofluorescence.

### Mitochondrial Profile Analysis

Mitochondrial profile assays were performed using Agilent Seahorse XF Cell Mito Stress Test Kit (Agilent Technologies, Inc., USA), in accordance with manufacturer’s instructions. Briefly, iPSCs or mDA neuron progenitors were seeded at an optimized density of 13 000 cells per well in an 8-well Seahorse cell culture plates coated with matrigel for iPSCs or polyornithine, laminine and fibronectine for mDA progenitors and incubated overnight. Each cell line was seeded in triplicates (N = 3). After 24 h, the Seahorse XFe Extracellular Flux Analyzer (Seahorse Biosciences, USA) was used to measure the OCR of each well. A period of 1 h before the measurements were initiated, the culture media in each well was replaced with 175 μL of Seahorse assay media supplemented with 1 mM pyruvate, and the plate was further incubated for 1 h at 37°C without CO2. Thereafter, successive OCR measurements was performed for each well, consisting of three basal OCR measurements, three OCR measurements following the automated injection of 1 μM oligomycin, three OCR measurements following the injection of 1 μM carbonyl cyanide p-trifluoromethoxyphenyl hydrazone (FCCP), and finally three OCR measurements following the dual injection of 1 μM rotenone and 1 μM antimycin A.

### JC-1 Assay

Cells in monolayer were suspended in 1ml warm medium. To the control cells 50 μM of CCCP was added and incubated at 37°C for 10 minutes. Tested cells were stained with 2 μM JC-1 and incubated at 37°C, 5% CO_2_ for 30 minutes in darkness. Then the cells were harvested with Accutase (BioLegend, USA) and washed with warm phosphate-buffered saline (PBS). Cell pellets were resuspended in warm culture medium and analyzed on a flow cytometer Attune Nxt Flow (Thermo Fisher Scientific).

### Cell surface marker screening by BD Lyoplate technology

The assay was processed as described by the manufacturer. iPS cells were detached from culture dishes by Accutase (BioLegend, USA), Next, 1×10^5^ cells were resuspended in 100 µl BPS per 96-well and stained in each well with the specific primary antibody for 30 min on ice. Then, the cells were washed and stained with the secondary antibody for 30 min on ice in darkness. Finally, the cells were resuspended in 250 µl PBS and collected (10.000 cells per well) by use of Attune NxT flow cytometer. The background fluorescence was compared with isotype controls. The analysis was performed by use of Attune NxT Software v2.2 and presented as a heat map generated in Excel 2021.

### Western Blots for OXPHOS complexes detection

Mitochondrial pellets were collected from HV and PD cell samples using Mitochondria Isolation Kit (Miltenyi Biotec, Germany). Mitochondrial proteins were extracted using Nuclear Extract Kit (Active Motif, Germany) according to manufacturer instructions. The protein concentration was determined by Bradford method. After SDS-PAGE and proteins transferred on PVDF membranes were incubated overnight at 4°C with Total OXPHOS Human WB Antibody Cocktail (Abcam, UK) and subsequently detected with HRP-conjugated goat anti-rabbit IgG secondary antibody (1:4000; Santa Cruz Biotechnology, USA). The membranes were developed with SuperSignal West Pico Chemiluminescence Substrate (Thermo Fisher Scientific, USA) and ChemiDoc MP Imaging System (Bio-Rad, USA) was used for membranes documentation.

### Patch clamp recording: data acquisition and analysis

Whole-cell recordings were performed on cultured iPSC at day 30-50 of differentiation. Neurons were plated on glass coverslips, transferred to a recording chamber, and constantly perfused (2 ml/min) with carbogenated, warm (32°C) ACSF composed of (in mM): 118 NaCl, 25 NaHCO3, 3 KCl, 1.2 NaH2PO4, 2 CaCl2, 1.3 MgSO4, and 10 glucose, pH 7.4, osmolality 290–300 mOsmol/kg. Glass micropipettes were fabricated from borosilicate glass capillaries (8-9 MΩ; Sutter Instruments) with the use of a horizontal puller (Sutter Instruments, Novato, CA, United States) and filled with a solution containing (in mM): 145 potassium gluconate, 2 MgCl2, 4 Na2ATP, 0.4 Na3GTP, 5 EGTA, 10 HEPES, pH 7.3 (osmolality 290-300 mOsmol/kg). The calculated liquid junction potential for these solutions was +15 mV and data were corrected for this value. Neurons were localized using an Examiner D1 microscope (Carl Zeiss) equipped with video enhanced infrared differential interference contrast. Cell-attached and subsequent whole-cell configurations were obtained under visual control using a negative pressure delivered by a mouth suction. Voltage- and current-clamp recordings were performed using a SEC 05LX amplifiers (NPI, Tamm, Germany), Micro 1401 mk II (Cambridge Electronic Design, Cambridge, United Kingdom), converter and Signal and Spike2 software (Cambridge Electronic Design). The recorded signal was low-pass filtered at 3 kHz and digitized at 20kHz. Action potential (AP) properties and time to first AP were calculated from the first AP evoked by a 500-ms-long depolarizing current pulse with an amplitude of + 90 pA. Excitability of recorded neurons was measured as the number of spikes in response to incremental (amplitude up to 140 pA, 10pA increment, 500 ms pulse duration) depolarizing current injections (input – output relationship, I-O). Second order polynomial curves were fitted to the I-O relationships and defined as neuronal gain. Additionally, AP threshold was assessed based on voltage response to current ramp (0 - 1 nA, 1 s) application. All used current stimuli were delivered from a membrane potential of − 75 mV, sustained with continuous current injections. Passive membrane properties (capacitance, resistance and time constant) were calculated and then averaged from the voltage response to 3 consecutive hyperpolarizing current pulses applied during zero current current-clamp recordings (−50 or - 100 pA, 1 s). The resting membrane potential and spontaneous AP were recorded in current clamp mode (zero holding current) for 3 min. Neuron was considered spontaneously active if at least one AP was generated during this time. Spontaneous postsynaptic currents were recorded in voltage clamp mode (− 50 mV holding voltage) for minimum of 3 min. Voltage-gated sodium channel blocker, tetrodotoxin (0.5 µM, Tocris Bioscience, Bristol, UK) and AMPA/NMDA receptors antagonists (10 µM CNQX and 50 µM DL-AP5, respectively, Sigma-Aldrich, Darmstadt, Germany) were used during the recordings. Custom MATLAB scripts (MathWorks Inc., Natick, MA, USA) were used for the final detection of electrophysiological parameters.

### Data preprocessing of microarray data – bioinformatics analysis

Microarray data were preprocessed using R/Bioconductor packages. Normalization was performed with three different approaches. The first one was done with limma package, using BGX file from Illumina site as reference. Data were normalized using the neqc() function (normexp background correction using negative control probes and quantile normalization using negative and positive control probes). Probe information was used for filtering. Probes with the quality ‘No match’ or ‘bad’ were removed. After normalization and filtering, 34476 probes were left. For the second and third approach, the beadarray package with illuminaHumanv4.db annotation was used. Subsequently, VSN or quantile normalization was performed. After normalization and filtering 34476 probes were left (the same as previously). To correct for possible batch effects, a surrogate variable analysis from the sva package was performed. For all three normalization approaches, all the data were used for analysis up to SVA. After SVA, the samples for KP and MM were filtered out from model matrix and data. Due to the fact that assessing the correct number of surrogate variables to account for, without disturbing downstream analysis, is complicated, we followed authors suggestions and two factors were accounted for.

### Differential gene expression analysis of microarray data – bioinformatic analysis

Limma package was used to perform differential expression analysis. Genes with a Benjamini-Hochberg corrected p-value lower than 0.05 were treated as differentially expressed.

### Functional analysis of microarray data – bioinformatic analysis

Gene Ontology analysis was performed using R/Bioconductor TopGO package. Terms were selected with parentChild algorithm with p-value cut-off set to 0.05.

### Statistical analysis

Statistical analysis of electrophysiological data was performed using GraphPad Prism for Windows (RRID: SCR_002798, GraphPad Software Inc., La Jolla, CA, United States). Normality of distribution was assessed, and data was analysed by Mann-Whitney and unpaired t-tests or Fisher’s exact tests where appropriate. All values are provided as mean ± SD (for normally distributed data) or median with interquartile range (for data that did not pass normality tests). Differences with a p value <0.05 were considered statistically significant.

## Results

### Generation of iPS cells from IPD and healthy volunteers

The group of healthy volunteers’ (HV) and PD patients consisted of participants with different gender and age. Detailed characteristics of the PD patients are described in Table S1. PD patients included in the study were without any mutations specific to Parkinson’s disease, which was analyzed by next-generation sequencing (NGS) (Figure 1).

**Figure 1.**
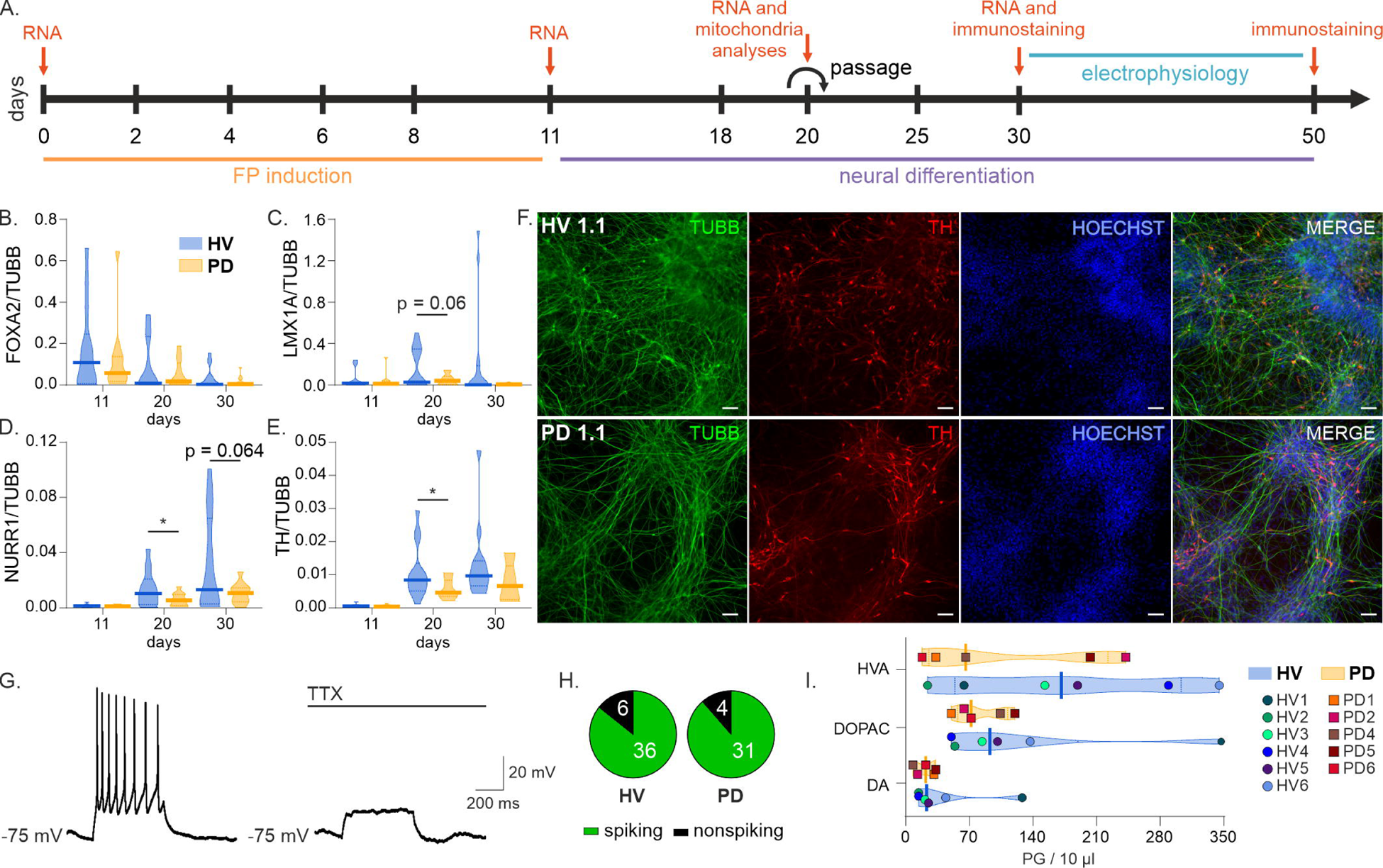
Differentiation of iPS cells into dopaminergic neurons. **(A)** Experimental procedure. **(B-F)** Relative expression of mDA neurons markers – FOXA2 **(B)**, LMX1A **(C)**, NURR1 **(D)** and tyrosine hydroxylase (TH) **(E)**. Medians ± IQRs presented. **(F)** Fluorescent microscopy photographs of immunohistochemical stainings against tubuline (green), TH (red) in healthy volunteer (top panel) and PD patient (lower panel). Nuclei marked with Hoechst staining (blue). **(G-H)** Majority of both, HV and PD cells was able to generate tetrodotoxin (TTX)-sensitive action potential (AP) upon depolarization with rectangular pulse **(G)**. No difference in occurrence of spiking and non-spiking neurons was observed **(H)** (Fisher’s exact test p = 0.748). **(I)** Extracellular concentrations of dopamine (DA) and its metabolites 3,4-Dihydroxyphenylacetic acid (DOPAC) and homovanillic acid (HVA). Medians ± IQRs presented.

Using Yamanaka’s factors, healthy volunteers’ (HV) and PD patients’ peripheral blood mononuclear cells (PBMCs) were reprogrammed and the obtained iPS cells were characterized. Analysis of expression levels of endogenous pluripotency markers (such as: NANOG, OCT3/4 and TERT), alkaline phosphatase staining, immunofluorescent staining of surface antigens (SSEA3, SSEA4, TRA1-60) and teratoma formation assay were performed (Figure S 1). All conducted tests confirmed the pluripotency of the derived iPS cell lines.

### Phenotype analysis of iPS cells

Using Lyoplate technology (BD) we evaluated the expression of 242 surface markers for four different PD iPS cell lines and four HV iPS cell lines. This study presents the first comprehensive phenotypic characterization of PD-derived iPS and comparative analysis of surface antigens expression for iPS cells for these two groups of donors. We identified 115 markers with expression levels ≥ 5%. The expression of each antigen was evaluated by analyzing the percent positive cells in the histograms compared to isotype control (not shown). The results for each sample were presented as a heat map generated in Excel 2021 (Figure S 2). Generally, we did not notice significant differences in the investigated panel of antigens between PD and HV groups. Variations in the antigen profile for the investigated iPS cell lines may be due to donor-specific variability.

### Differentiation of iPS cells into dopaminergic neurons

Subsequently, iPS cells were differentiated to dopaminergic neuron using Kriks et al. protocol (Fig. 1 A) ^15^. During the differentiation process, cells were collected at each stage of differentiation. FOXA2, LMX1A, NURR and TH genes were differentially expressed in differentiating HV and PD cells (Fig. 1 B - E). On day 30, TH and TUBB staining was performed, and we observed similar levels of TH and TUBB positive cells in HV and PD patients ‘s neurons cells were at (Fig. 1 F). Differences in morphology of progenitor cells and mature neurons were not observed between groups (Figure S2 A)

To examine the electrophysiological properties of cultured neurons, 42 cells from six HV and 35 cells from five PD patients were recorded in whole-cell patch-clamp mode. The ability to generate action potentials (APs) in response to depolarizing current pulses was tested to assess whether the examined cells are mature neurons. Applied depolarizing current pulses evoked action potentials in 85.7% (36 out of 42) HV and 88.57% (31 out of 35) of PD neurons (Fig. 1 G, H), indicating that most differentiated cells were functional. Ability to generate APs did not differ between cells from HV and PD patients (p = 0.748, Fisher’s exact test) (Fig. 1 H). APs generated by both HV and PD patient’s neurons were sensitive to voltage-gated sodium channel blocker tetrodotoxin (TTX, Fig. 1 G right panel).

Extracellular concentration of DA, 5-HT, DOPAC, HVA, and 5-HIAA, measured using a high-performance liquid chromatography (HPLC) with electrochemical detection, were also similar in HV and PD samples (Fig. 1 I).

### Mitochondrial status in IPD

In TEM analysis we observed numerous, small (∼[0.7 to 2.5 µm), spherical shape mitochondria in cytoplasm of the HV and PD iPS cells (Fig. 2 A). Mitochondria had typical morphologic features with internal electron-dense cristae. Comparative TEM analysis showed no significant differences that could be associated with mitochondrial dysfunction either in ultrastructure or in number of these organelles between PD and HV cells.

**Figure 2.**
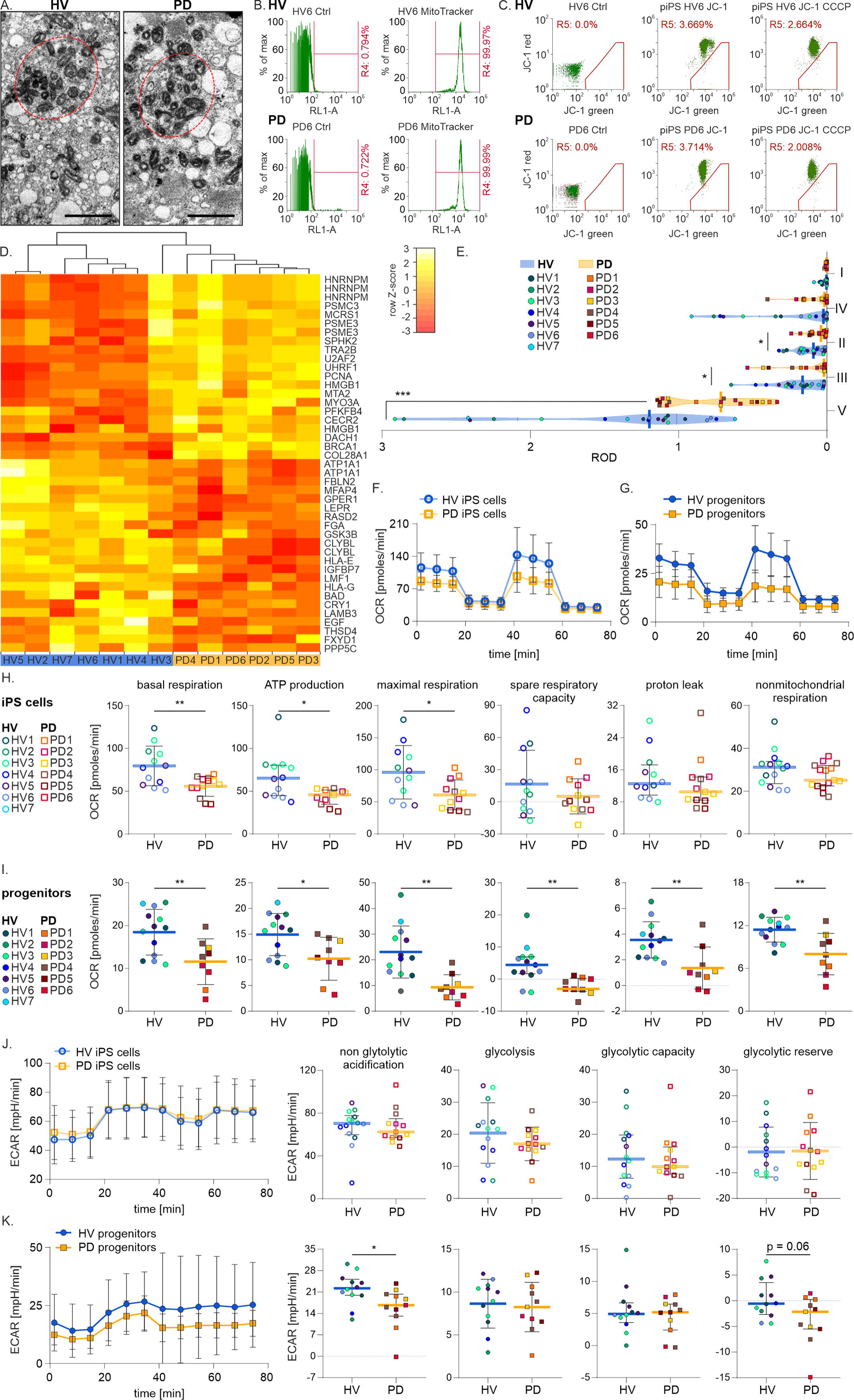
Mitochondria status in iPS cells and iPSC-derived mDA progenitors. **(A)** A comparative transmission electron microscopy images of iPS HV and PD mitochondria; Scale bar = 5 µm. **(B)** iPS cells untreated (left) and stained with MitoTracker Red (right). **(C)** Cytofluorimetric analysis of mitochondrial membrane potential using JC-1 MitoProbe. **(D)** Heatmap for transcripts corresponding to genes assigned to pre-selected GO terms and classified as significantly differentially expressed by all 3 methods. **(E)** WB results of mitochondrial complexes protein levels in heathy volunteers (HV) and Parkinson’s (PD). Quantitative results presented on violin plots are based on densitometry of the analyzed complexes for HV and PD respectively. Medians ± IQRs presented. **(F-K)** Seahorse Extracellular Flux Assay. (**F-I)** Profile of oxygen consumption rate (OCR) of patient-derived iPS and patient-derived neuronal progenitors. **(J-K)** Profile of the extracellular acidification rate (ECAR) of patient-derived iPS and patient-derived neuronal progenitors.

Cytometric assessment of mitochondria using mitochondrial-specific dye (MitoTracker) showed highly positive signal for both HV and PD patient’s (Fig. 2 B), but expression levels were similar and reached over 99%. We also noticed no significant variations in MFI level (not shown). Flow cytometry analysis of mitochondria membrane potential using JC-1 MitoProbe showed similar levels of plasma membrane depolarization have been detected in HV and PD samples (Fig. 2 C). These results indicate a lack of significant differences in mitochondrial mass or basic mitochondrial fitness between PD and HV.

Transcriptome analysis revealed genes that were differentially expressed between progenitors from healthy volunteers’ and PD patients’ (Fig. 2 D). These results and reports from the literature describing mitochondrial defects found in PD ^16^ allowed us to compare the results of the transcriptome analysis with the mitochondrial proteome. We prepared a list of transcripts that were translated, and their levels in mitochondria differed between a group of healthy volunteers’ and PD patients’.

Since we did not observe variations in mitochondria mass or mitochondrial activity, we analyzed the mitochondrial OXPHOS complexes (I to V) level in HV and PD derived cells. Indeed, we found significant differences at protein levels in complex II, III and V between samples from healthy donors and PD patients (Fig. 2 E).

### Mitochondria respiration status in iPS cells and iPSC-derived mDA progenitors

Next, we determined the oxygen consumption rates (OCR) and extracellular acidification rate (ECAR) by iPS and iPS-derived mDA progenitors derived from HV and PD. We found a tendency of decreased OCR and ECAR in iPSC and iPSC-derived mDA progenitors from PD patients compared to HV control (Fig. 2 F, G). Basal respiration was significantly lower in PD than HV both in iPS and iPS-derived progenitors (Fig. 2 H). Spare respiration was also significantly lower in iPS-derived progenitors in PD than in HV (practically there was no spare respiratory capacity in PD derived cells) (Fig. 2 I). ATP-linked respiration was significantly lower in PD than HV both in iPS and iPS-derived progenitors (Fig. 2 H, I). Moreover iPS-derived mDA progenitors from PD patients showed highly significant difference between oligomycin and rotenone inhibition of OCR, representing energy substrate (ATP) independent respiration and an increase in the passive leak of protons from the inner mitochondrial membrane (Fig. 2 I). Those results correspond to increased acidification in iPSC-derived mDA progenitors visible in ECAR (Fig. 2 K). Maximal respiration was significantly lower in iPSC-derived mDA progenitors in PD patients than HV similarly as in iPSC (Fig. 2 K). In iPSC-derived mDA progenitors non-mitochondrial oxygen consumption was significantly lower in PD group than in HV, whereas in iPSC there was no such a difference between groups (Fig. 2 J, K).

### Neurophysiological status of IPD neurons

Whole-cell patch-clamp recordings revealed that spontaneous resting membrane potential of HV neurons was significantly more hyperpolarized in comparison to PD neurons (p = 0.02, unpaired t-test) (Fig 3 A, top left and bottom right panels; Table 1). Moreover, spontaneous activity of studied neurons was examined in zero holding current mode. In HV neurons, we observed spontaneously generated APs in 46.88% (15 out of 32) of recorded cells, whereas in PD neurons 30.43% (5 out of 21) of cells spontaneously generated APs (p = 0.147, Fisher’s exact test) (Fig. 3 A, bottom left panel). Frequency of APs generated by spontaneously active neurons did not differ between groups (p = 0.22, Mann-Whitney test) (Fig. 3 A, top right panel; Table 1).

**Figure 3.**
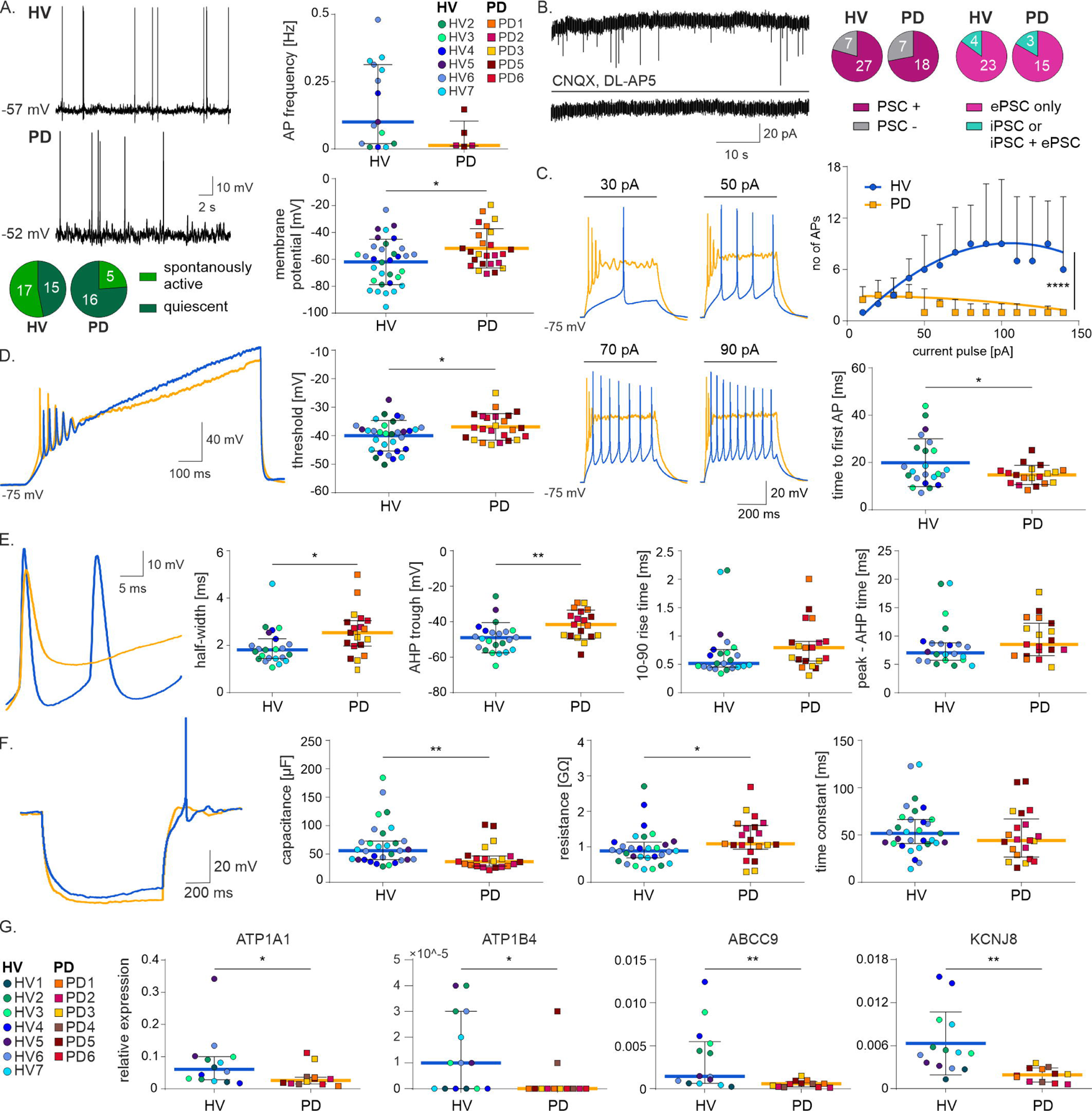
Neurophysiological status of IPD neurons. **(A)** Top and middle left panels: traces of zero current current-clamp recordings of spontaneously active HV (top) and PD (middle) neurons. Bottom left panel: Occurrence of spontaneously active neurons within groups. Top right panel: frequency of spontaneously generated APs. Medians ± IQR presented. Bottom right panel: Resting membrane potential (t test, p = 0.02). Mean ± SD presented. **(B)** Top left panel: voltage clamp recording (holding potential −50 mV) in sACSF. Excitatory postsynaptic currents (ePSC) visible as inward current. Recorded ePSC vanish in the presence of AMPA/NMDA receptors antagonists (CNQX and DL-AP5) (bottom left panel). Middle panel: Occurrence of postsynaptic currents (PSC). Both in HV and PD, in majority of neurons PSC were recorded (PSC + neurons). Right panel: Nature of recorded postsynaptic currents. Only minority of neurons exhibited inhibitory postsynaptic currents (iPSC). **(C)** Left panel: Depolarizing current pulses of increasing intensity induce increasing number of generated APs in HV, but not in PD neurons. Top right panel: PD neurons exhibit significantly lower excitability that HV neurons (comparison of fitted second order polynomial curves, F (3,22) = 611.2, p < 0.0001. Median ± IQR presented. Bottom right panel: Concomitantly with lower excitability, PD neurons display shorter time to first generated AP, when depolarized using +90 pA current pulse (t test, p = 0.043). Mean ± SD presented. **(D)** Left panel: Voltage response do depolarizing current ramp. Right panel: Threshold for generation of AP is more depolarized in PD neurons (t test, p = 0.028). Mean ± SD presented. **(E)** Leftmost panel: First AP generated by HV and PD neuron upon depolarization with +90 pA current pulse. Left panel: half-with of single AP (Mann-Whitney, p = 0.018). Median ± IQR presented. Middle panel: AHP trough of recorded neurons (t test, p = 0.006). Mean ± SD presented. Righ and rightmost panels: 10-90 rise and peak-AHP time displayed by recorded neurons, respectively. Median ± IQR presented. **(F)** Leftmost panel: voltage response to the hyperpolarizing (−50 pA) current pulse during zero current current-clamp recording. Left panel: electrical capacitance of recorded neurons (Mann-Whitney, p = 0.004). Median ± IQR presented. Right panel: Neuronal membrane’s resistance (Mann-Whitney, p = 0.031). Median ± IQR presented. Rightmost panel: Membrane’s time constant doesn’t differ between groups. Median ± IQR presented. **(G)** ATP1A1, ATP1B4, ABCC9 and KCNJ8 levels in differentiated iPS cells obtained from HV and PD patients. qPCR results were calculated with the ΔCt method, and GAPDH served as a constitutive control. Median ± IQR presented.

**Table 1.** Electrophysiological properties of HV and PD neurons. ° unpaired t-test; Mann – Whitney unpaired test; bold values denote statistical significance at the p < 0.05 level.

|  | HV | PD | statistics |
| --- | --- | --- | --- |
| **membrane properties** |  |  |  |
| resting membrane potential [mV] | $-61.88 \pm 16.93$ | $-51.8 \text{ mV} \pm 14.59$ | ° **$p = 0.02$** |
| time constant [s] | $51.72 \pm 25.32$ | $44.18 \pm 40.3$ | □ $p = 0.323$ |
| resistance [GΩ] | $0.89 \pm 0.44$ | $1.08 \pm 0.67$ | □ **$p = 0.031$** |
| capacitance [μF] | $55.63 \pm 32.67$ | $36.31 \pm 17.85$ | □ **$p = 0.004$** |
| **firing properties** |  |  |  |
| spontaneous firing rate [Hz] | $0.1 \pm 0.29$ | $0.013 \pm 0.092$ | □ $p = 0.22$ |
| time to first AP [ms] | $19.94 \pm 10.10$ | $14.77 \pm 4.06$ | ° **$p = 0.043$** |
| threshold [mV] | - 39.99 ± 5.36 | - 36.89 ± 4.67 | ° **p = 0.028** |
| **single AP properties** |  |  |  |
| threshold [mV] | -39.07 ± 4.86 | -36.45 ± 3.77 | ° p = 0.063 |
| 10 – 90 rise [μs] | 19.94 ± 10.10 | 14.77 ± 4.06 | ° **p = 0.043** |
| half-width [ms] | 1.81 ± 0.83 | 2.54 ± 1.08 | □ **p = 0.018** |
| peak – AHP time [ms] | 7.04 ± 3.06 | 8.53 ± 5.69 | □ p = 0.164 |
| AHP trough [mV] | -49.0 ± 8.52 | -41.55 ± 8.15 | ° **p = 0.006** |

To verify whether HV and PD neurons are functionally connected, we performed voltage-clamp recordings (holding potential – 50 mV) (Fig. 3 B, left panel). Spontaneous postsynaptic currents were recorded from 79.41% (27 out of 34) HV neurons and 72% (18 out of 25) PD patients’ neurons, and there was no difference in the ratio of their occurrence between studied groups (p = 0.549, Fisher’s exact test) (Fig 3 B, middle panel). Majority of recorded neurons was shown to receive excitatory input, as in 23 out of 27 recorded HV neurons only excitatory postsynaptic currents (ePSCs) were observed (in remaining three both excitatory and inhibitory postsynaptic currents (iPSCs) were recorded, and in one neuron only iPSCs were observed) (Fig. 3 B, right panel). Similarly, in 15 PD (out of 18 exhibiting PSC) neurons, only ePSCs were observed, and in the remaining three, both ePSCs and iPSCs were visible (Fig. 3 B, right panel). Recorded ePSCs were AMPA/NMDA receptors dependent, as they vanished upon ACSF enrichment with those receptors’ antagonists – CNQX and DL-AP5 (Fig. 3 B, bottom left panel).

To test possible differences in excitability of HVs’ and PD patients’ neurons, we used depolarizing current steps (10 pA increment) to calculate neuronal gain from the number of APs evoked in response to the current steps (Fig. 3 C, left panel). Neuronal gain was significantly different between neurons from HV and PD patients, with PD neurons exhibiting lower increase in number of generated APs in response to consecutive pulses in comparison to HV (comparison of fitted second order polynomial curves, F (3,22) = 611.2, p < 0.0001) (Fig. 3 C, top right panel, Table 1). Moreover, time from the beginning of the pulse to the first generated AP was significantly shorter in neurons from PD patients (p = 0.043, unpaired t-test) (Fig 3 C, right bottom panel, Table 1). Additionally, ramp current injections revealed a significant difference in APs threshold in examined neurons (Fig. 3 D, left panel), with more depolarized value recorded from PD neurons when compared to HV cells (p = 0.028, unpaired t-test) (Fig. 3 D, right panel; Table 1).

Analysis of the action potential shape evoked by depolarizing current pulse (+90 pA) (Fig. 3 E, leftmost panel) revealed significant differences between HV and PD neurons, with the latter having broader half-width (Fig. 3 E, left panel; Table 1) and more depolarized AHP trough (Fig. 3 E, middle panel; Table 1). No differences were observed in other analyzed parameters of AP shape (Fig. 3 E, right and rightmost panels; Table 1).

To assess possible differences in passive membrane properties of HV and PD neurons, we applied hyperpolarizing current steps (−50 or −100 pA depending on a neuron, to hyperpolarize it to at least −80 mV) during current-clamp (zero holding current) recordings (Fig. 3 F, leftmost panel). Neurons from PD patients had significantly smaller electrical capacitance and higher resistance when compared to healthy volunteers’, with no difference in the time constant (Fig. 3 F, left, right and rightmost panels, respectively; Table 1).

Finally, we analyzed mRNA expression levels of ATP Binding Cassette Subfamily C Member 8 and 9 (ABCC8, ABCC9), Potassium Inwardly Rectifying Channel Subfamily J Member 8 and 11 (KCNJ8, KCNJ11) and ATPase Na+/K+ Transporting Subunits Alpha 1-4 and Beta 1-4 (ATP1A1-4 and ATP1B1-4) in differentiated iPS cells obtained from HV and PD patients. Importantly, differentiated PD cells displayed diminished levels of ATP1A1, ATP1B4, ABCC9, KCNJ8 (Fig. 3 G).

## Discussion

Parkinson disease is caused by well-studied and characterized inherited genetic mutations and by to large extends unknown environmental causes, in both cases involving atypical accumulation of α-synuclein in Levy bodies ^8–10^. Since IPD accounts for approximately 90% of all PD cases and in contrast to genetic form of PD its etiology is mostly unknown, we focused on dissecting the biological mechanisms responsible for this form of PD.

We did not observe major differences in differentiation potential between IPD and HV derived DA neurons, which is contrary to previously published studies that observed changes in biology of DA neurons including proliferation and differentiation ^17^. However, these studies were carried on predominantly using iPS cells derived from familial form of PD ^18^ or YOPD ^13^, whereas we used IPD derived iPS cells. Accordingly, we did not observe statistically significant differences in production of DA and its metabolite derivatives, whereas previous studies suggested different DA production levels in neurons derived from iPS cells ^19^. Additionally, we found that the ability to generate both spontaneous and evoked APs did not differ between cells from HV and PD patients. The differences in the origin of neurons used in the current and cited studies most probably underlie the observed differences in DA neuron biology, highlighting the importance of studying idiopathic PD as a distinct entity with unique features and characteristics.

Since mitochondrial dysfunctions related to genetic aberration have been described in the course of Parkinson’s disease, we looked at different aspects of mitochondria biology. Importantly, we observed differences in the mitochondria respiratory status, whereas there were no differences in mitochondria mass, morphology or membrane potential in cells derived from HV and PD.

OXPHOS mitochondrial complexes were differentially expressed between HV and PD groups, which resulted in changes in mitochondria respiratory status. Since the OXPHOS process generates energy in mitochondria in the form of ATP ^20^ the disruption of mitochondrial function may hinder the production of ATP and disrupt the functioning of dopaminergic neurons ^16, 21^.

Furthermore, our results suggest also differences in ATP turnover. We observed lower expression levels of Na^+^/K^+^-ATPase subunits. This ubiquitous protein complex is responsible for the generation and maintenance of the Na^+^/K^+^ gradient across the cell membrane and plays a major role in generating the voltage across the membrane and is critical for the proper cell functioning ^22^. Importantly, our observations indicate that PD neurons have a more depolarized membrane potential when compared to HD neurons, which may be attributed to the lower expression of Na+/K+-ATPase ^23^. Notably, mutations in this complex have been also associated with early onset of PD ^24–26^. Interestingly, α-synuclein has been shown to block and disrupt the function of Na^+^/K^+^-ATPase, resulting in an increase in the intracellular Na^+^ concentration what may lead to attenuation of Ca^2+^ extrusion via the Na^+^/Ca^2+^ exchanger and neuronal cell death ^27, 28^. Reduction of Na^+^/K^+^-ATPase activity has been also linked to aggravation of α-synuclein-induced pathology, including a reduction in tyrosine hydroxylase (TH) activity and motor disfunctions ^29^. Furthermore, recent data suggest also that Na^+^/K^+^-ATPase expression and activity is regulated by Leucine-Rich Repeat Kinase 2 strongly implicated in the pathophysiology of Parkinson’s disease ^30^.

Our transcriptomic analysis revealed also different expression levels of ATP-sensitive K^+^ (K-ATP) channel subunits. We observed lower expression of potassium inwardly rectifying channel member - Kir6.1 (KCNJ8) and ATP binding cassette member - Sur2 (ABCC9). Diminished levels of KCNJ8 and ABCC9 may result in decreased K-ATP channel activity and reduced hyperpolarization of the membrane potential ^31^, which could contribute to the observed in the current studies, differences in resting membrane potential between HV and PD neurons. Activity of ATP-sensitive K^+^ channels play an important role in regulating the duration and shape of the action potential by contributing to its repolarization phase ^32, 33^, therefore observed decreased expression of mRNA encoding K-ATP channel subunits may underly observed prolonged duration of the action potential in PD neurons and less hyperpolarized value of AHP trough. Importantly, changes in action potential shape, and in particular, impaired recovery of membrane polarity after action potential generation, may underlie the observed lower excitability of PD neurons. Neuronal excitability in animal models od PD was shown to be reduced due to disrupted ionic mechanisms for AP generation ^34^ and diminished dopaminergic input to the motor circuits ^35, 36^, and in neurons derived from familial Parkinson’s disease patients ^37^. At the same time, no changes in excitability, were reported for neurons derived from iPD patients ^37^, however, in cited study only the total number of action potentials generated in response to current inputs was counted, and changes in response to varying current pulse strength was not investigated. Therefore, to the best of our knowledge, our data are the first showing impairments in neuronal gain in DA neurons derived from iPD patients, what translates into their impaired ability to correctly respond to the input signals. Impairment in neuronal gain in DA neurons derived from iPD, as observed in our study, can lead to dysfunctional circuit activity and contribute to the symptoms of Parkinson’s disease.

Mutations in both KCNJ8 and ABCC9 genes have been previously linked to neurological diseases or disorders, such as hippocampal sclerosis, depression, and sleep disorder ^38–42^. Interestingly, dysfunction of K-ATP channel due to mutations of ABCC9 may lead to neurodegeneration ^41, 42^. Furthermore, upregulation of another ATP binding cassette member – Sur1 (ABCC8) correlated with neuronal damage in transgenic mice expressing A53T mutant α -synuclein ^43^. In chronic mouse models of Parkinson’s disease, K-ATP channels might be involved in neurodegeneration of DA neurons, since activation of another ATP binding cassette member - Kir6.2 (KCNJ11) led to degeneration of midbrain DA neurons ^44^. Similarly, elevated mRNA expression of Kir6.2 and a high in vivo bursting was observed in human substantia nigra DA neurons from Parkinson’s disease patients ^45^.

Importantly, we observed also lower OCR (oxygen consumption rate) and ECAR (extracellular acidification rate) values in mDA progenitors derived from iPSCs and iPSCs from PD patients compared to the HV control group. Similarly, other values based on OCR and ECAR measurement such as basal respiration, spare respiration, and ATP-related respiration were also significantly lower in PD than HV in both iPSC and iPSC-derived progenitors. Importantly, observed decrease in the availability of ATP and in the energy consumption, as well as changes in acidification, may constitute contributing factors to the observed reduced neuronal excitability of PD patients DA neurons ^46–50^.

In addition, mDA derived from iPSCs progenitors from PD patients showed a highly significant difference between oligomycin and rotenone inhibition of OCR, representing ATP independent respiration and an increase in passive proton leakage from the inner mitochondrial membrane. These results correspond to the increased acidification in iPSC-derived mDA progenitors seen in ECAR. Maximal respiration was significantly lower in iPSC-derived mDA precursors in PD patients than HV similar to iPSC. In mDA progenitors derived from non-mitochondrial iPSCs oxygen consumption was significantly lower in the PD group than in the HV group, while in the iPSC group there was no difference between the groups. The mechanisms underlying the low OCR (oxygen consumption rate) in Parkinson’s disease are not yet fully understood, but several theories have been proposed. One theory points to mitochondrial dysfunction, which may result in decreased OCR. Mitochondrial dysfunction can lead to decreased ATP production and increased oxidative stress, which can further disrupt mitochondrial function and contribute to disease progression ^51, 52^. Furthermore, dopamine has been shown to regulate mitochondrial function, and a decrease in dopamine levels may result in decreased OCR and contribute to disease progression ^51, 53, 54^. Parkinson’s disease is also associated with increased oxidative stress, which can damage cellular components and disrupt cellular processes, including mitochondrial function ^55^. Increased oxidative stress may result in decreased OCR and contribute to disease progression. Chronic inflammation is also associated with Parkinson’s disease and can lead to decreased OCR by disrupting mitochondrial function and increasing oxidative stress ^10, 16, 21, 49^. This can also contribute to the progression of the disease. In summary, the mechanisms underlying low OCR in Parkinson’s disease are complex and multifactorial, involving mitochondrial dysfunction, dopamine deficiency, oxidative stress, and inflammation ^54^. Despite overwhelming evidence suggesting that mitochondrial function is important for the pathogenesis of PD, the ability to identify the initial event in the cascade of changes that lead to neurodegeneration remains elusive ^56^.

## Acknowledgments

We would like to thank O. Woźnicka from the Department of Cell Biology and Imaging, Institute of Zoology and Biomedical Research, Jagiellonian University, for her skilled technical assistance with TEM material preparation and observations.

