## Supplementary Figure 1 for "Mitochondrial fitness influences neuronal excitability of dopaminergic neurons from patients with idiopathic form of Parkinson’s disease"

(A)

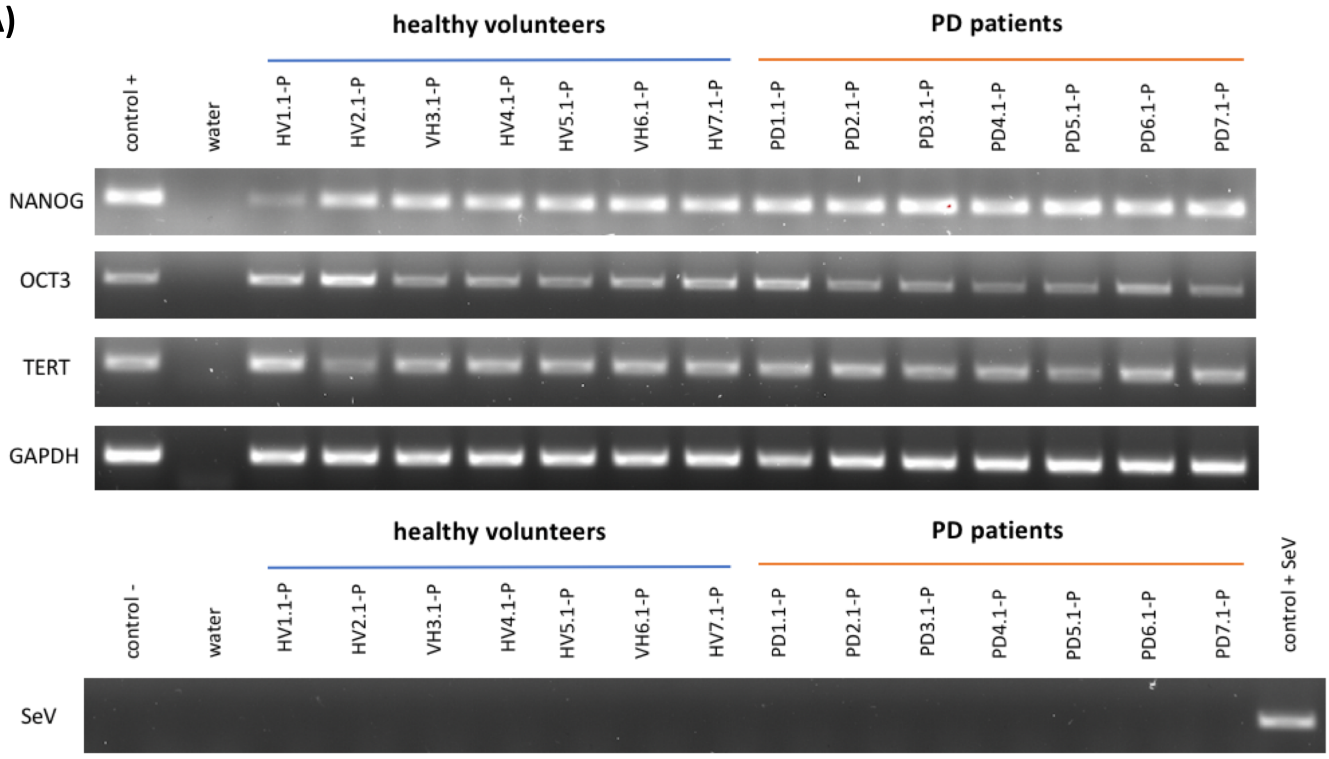


(B)

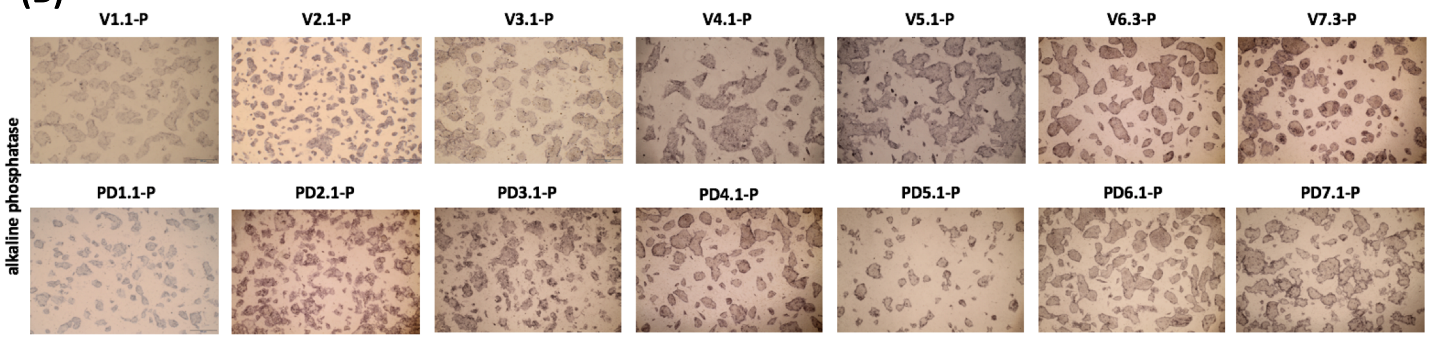


(C)

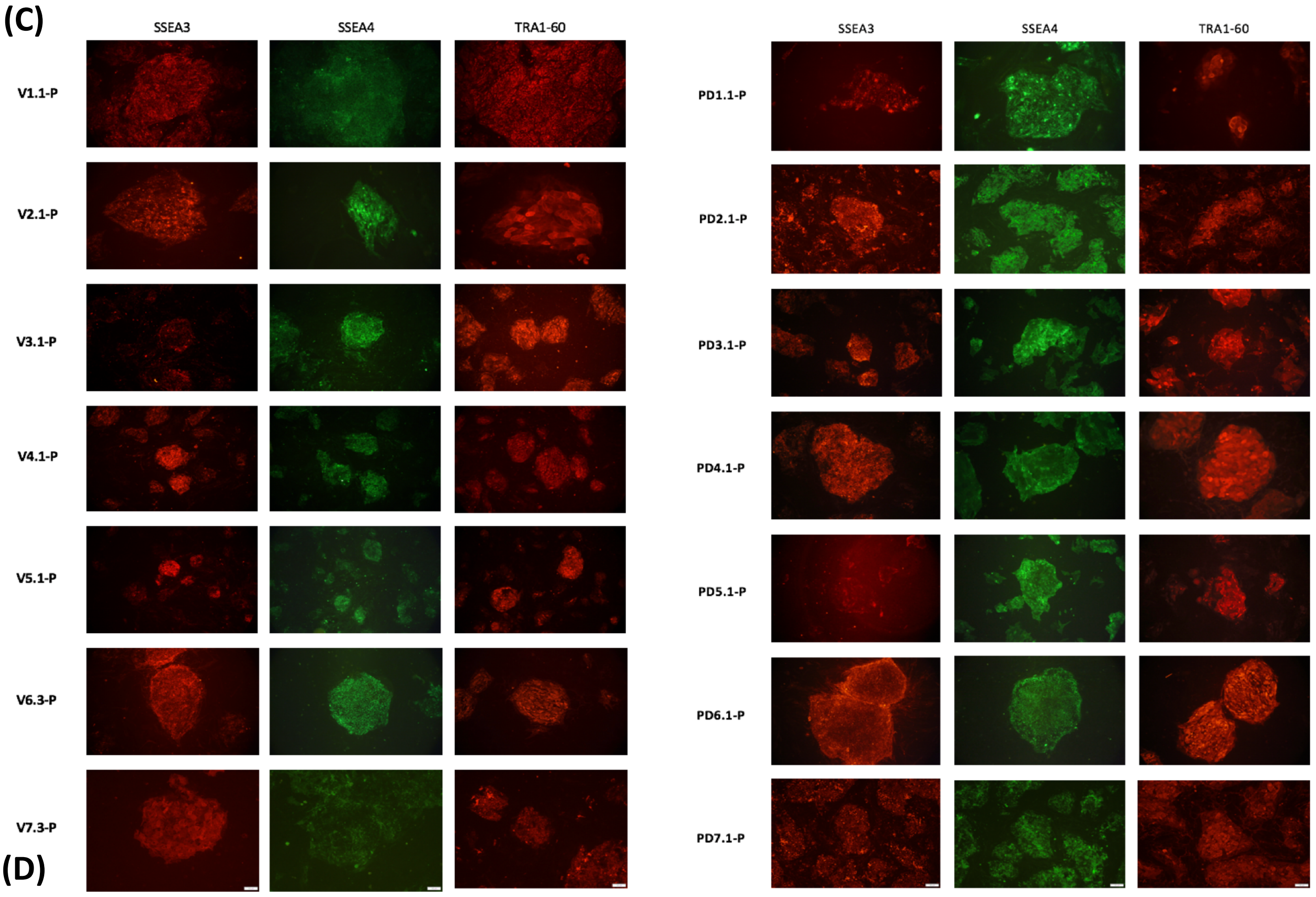


(D)

| cell lines / three germ layers | V1.1-P | V2.1-P | V3.1-P | V4.1-P | V5.1-P | V6.3-P | V7.3-P | PD1.1-P | PD2.1-P | PD3.1-P | PD4.1-P | PD5.1-P | PD6.1-P | PD7.1-P |
|---|---|---|---|---|---|---|---|---|---|---|---|---|---|---|
| ectoderm | + | + | + | + | + | + | + | + | + | + | + | + | + | + |
| endoderm | + | + | + | + | + | + | + | + | + | + | + | + | + | + |
| mesoderm | + | + | + | + | + | + | + | + | + | + | + | + | + | + |

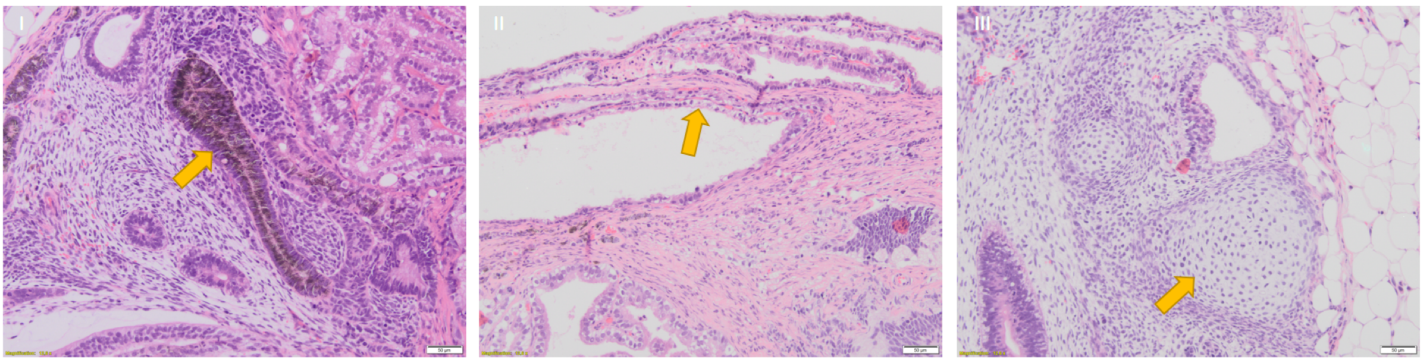

**Fig. 2| Characterization of generated iPS cell lines from seven healthy volunteers and seven Parkinson's disease (PD) patients. a**, NANOG, OCT3/4 and TERT (endogenous pluripotency markers) expression in iPS cell lines after $10^{th}$ passage by RT-PCR. GAPDH is a housekeeping control. At $10^{th}$ passage the Sendai virus transgen (SeV) were not detected. piPS (commercially available protein induced iPS cell line) was a positive control for endogenous pluripotency markers and negative control for SeV transgen. sv-iPS - iPS cell line on early passage with SeV transgene, used as positive control. $H_2O$ was used as a control of reaction. **b**, Images of iPS cells cultures showing alkaline phosphatase (AP) staining. **c**, Immunofluorescent staining of surface antigens (SSEA3, SSEA4, TRA1-60) of generated iPS cell lines. **d**, Teratomas were generated by iPS cell lines in vivo. Histological analysis shown tissues characteristic for all three germ layers. Representative images of these structures: pigmented cells (I), secretory epithelium (II), cartilage (III). NANOG—homebox protein NANOG; OCT3/4—octamer binding transcription factor 3/4; TERT—telomerase; GAPDH—housekeeping gene, glyceraldehyde 3-phosphate dehydrogenase; SEV—primer specific for Sendai Virus genome; SSEA3—stage specific embryonic antigen 3; SSEA4—stage specific embryonic antigen 4; TRA1-60—podocalyxin. Similar results were observed in other clones.
