## Supplementary figures and images for "Mitochondrial fitness influences neuronal excitability of dopaminergic neurons from patients with idiopathic form of Parkinson’s disease"

### Supplementary Figure 2

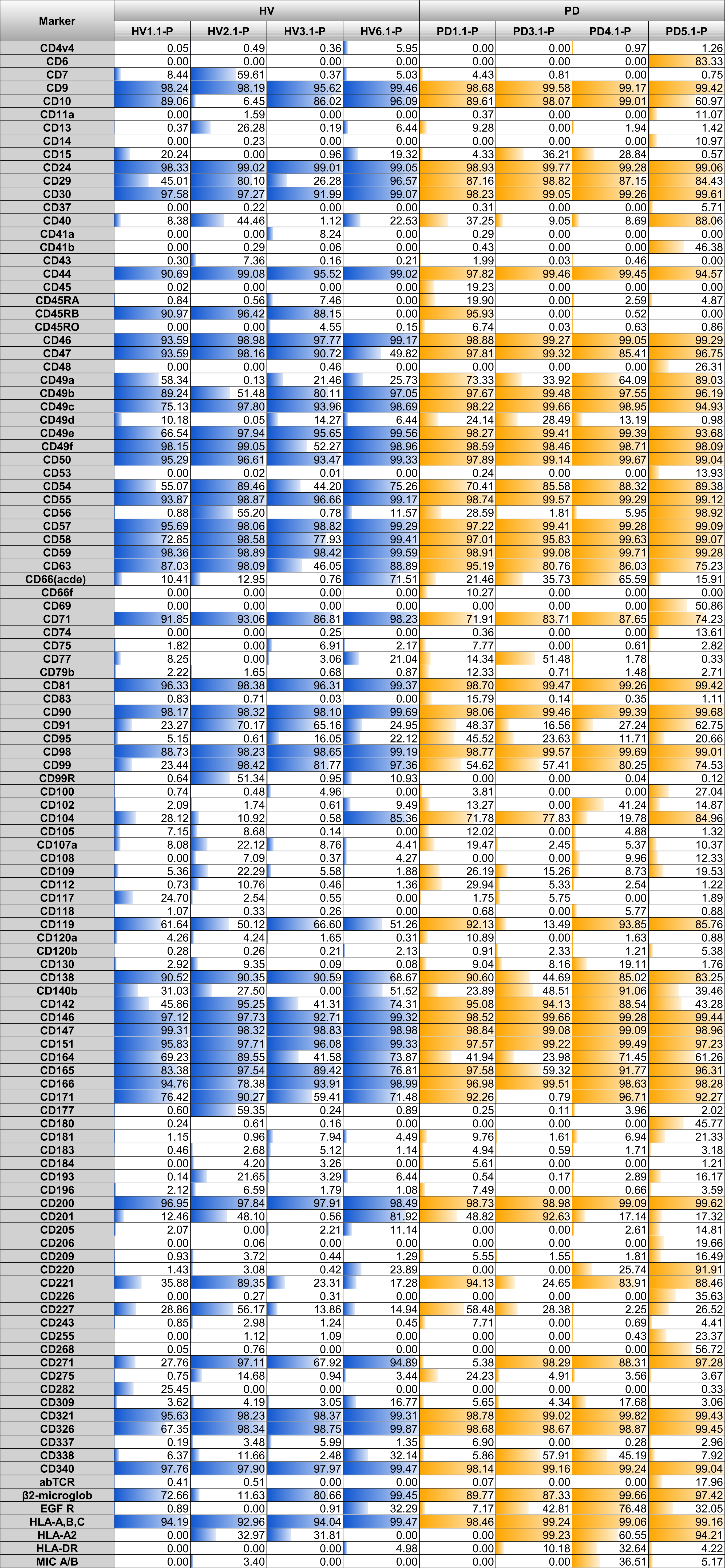

### Supplementary Table 1

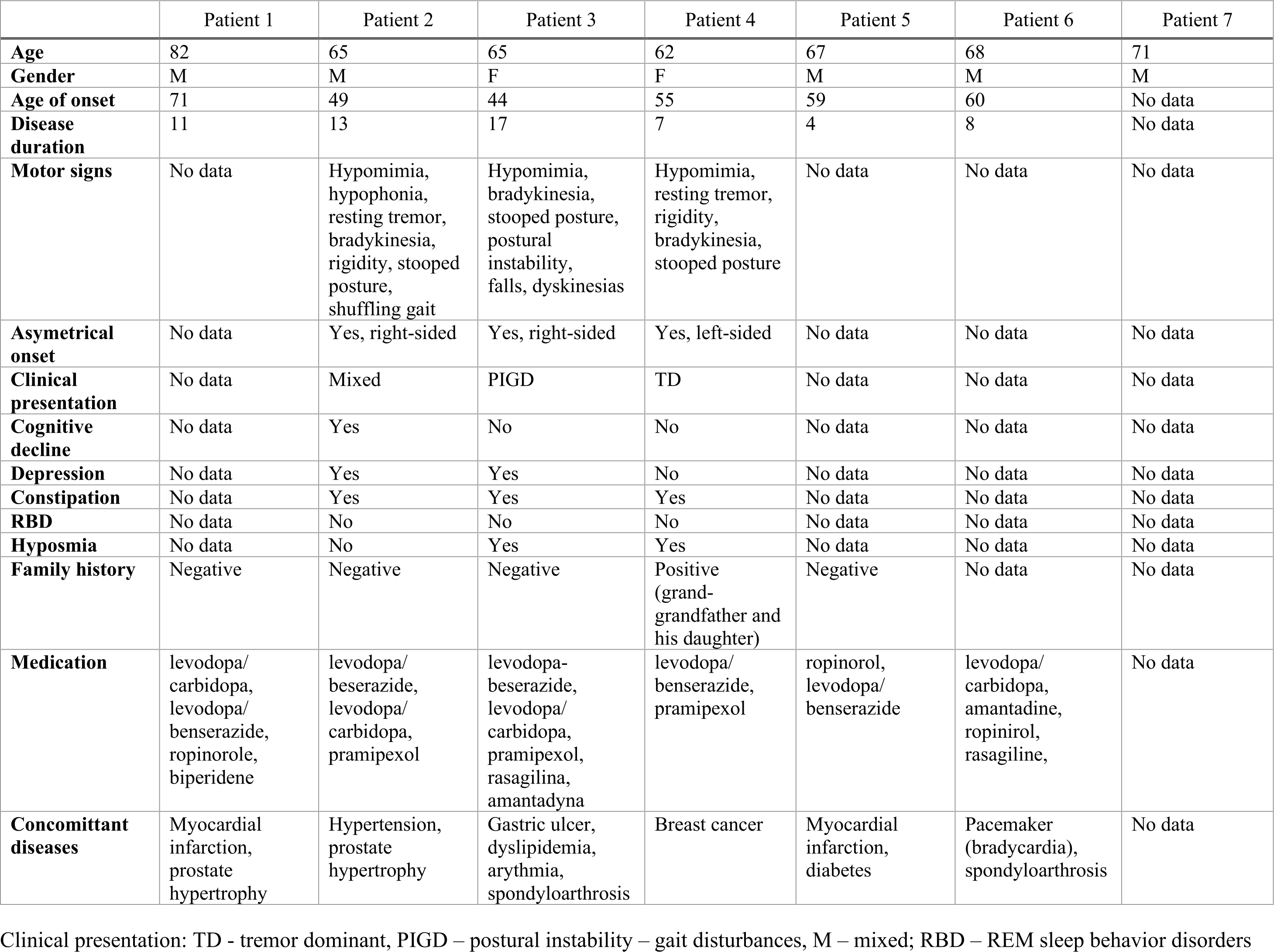
